# Improving a probabilistic cytoarchitectonic atlas of auditory cortex using a novel method for inter-individual alignment

**DOI:** 10.1101/2020.03.30.015313

**Authors:** Omer Faruk Gulban, Rainer Goebel, Michelle Moerel, Daniel Zachlod, Hartmut Mohlberg, Katrin Amunts, Federico De Martino

**Affiliations:** Department of Cognitive Neuroscience, Maastricht University, The Netherlands; Brain Innovation B.V., Maastricht, The Netherlands; Maastricht Centre for Systems Biology, Faculty of Science and Engineering, Maastricht University, The Netherlands; Institute for Neuroscience and Medicine (INM-1), and JARA Brain, Research Centre Jülich, Jülich, Germany; C. and O. Vogt Institute for Brain Research, Heinrich Heine University Düsseldorf, Germany; Center for Magnetic Resonance Research, University of Minnesota, Minneapolis, MN, USA

## Abstract

The human superior temporal plane, the site of the auditory cortex, displays a high inter-individual macro-anatomical variation. This questions the validity of curvature based alignment (CBA) methods for in vivo imaging data. Here, we have addressed this issue by developing CBA+, which is a cortical surface registration method that uses prior macro-anatomical knowledge. We validate this method by using cyto-architectonic areas on ten individual brains (which we make publicly available). Compared to volumetric and standard surface registration, CBA+ results in a more accurate cyto-architectonic auditory atlas. The improved correspondence of micro-anatomy following the improved alignment of macro-anatomy validates the superiority of CBA+ compared to CBA. In addition, we use CBA+ to align in vivo and post mortem data. This allows projection of functional and anatomical information collected in vivo onto the cyto-architectonic areas, which has potential to contribute to ongoing debate on the parcellation of the human auditory cortex.

## Introduction

Historically, there has been a substantial effort to describe the micro-anatomy of the human auditory cortex (***Von Economo and Horn, 1930; Galaburda and Sanides, 1980; Rivier and Clarke, 1997; Morosan et al., 2001; Wallace et al., 2002; Morosan et al., 2005; Clarke and Morosan, 2012; Nieuwenhuys, 2013***). Various parcellation schemes have been proposed, which identify a primary area (core; primary auditory cortex) as well as secondary belt and tertiary parabelt auditory areas (***Rivier and Clarke, 1997; Moerel et al., 2014***). The primary auditory cortex (PAC) is generally located on the medial two-thirds of Heschl’s Gyrus.

It has proven challenging to use these results to identify auditory areas in individuals in vivo, as classical cyto- (and myelo-) architectural approaches are limited by the absence of an objective metric defining cyto-architectonic areas. In addition, relating micro-anatomical characteristics to macro-anatomy is hampered by the inherent two-dimensional representation of the results (i.e. by means of drawings or labelled slices) and scarce information regarding inter-subject variability. Instead, observer independent methods for the analysis of serial cyto-architectonically stained sections, that additionally correct for shrinkage artifacts typical of histological processing (***Amunts et al., 2000***), have been developed in the last 20 years (***Schleicher et al., 1999***). Using this method, ***Morosan et al. (2001***) identified various auditory areas in the superior temporal cortex and generated a probabilistic atlas based on ten individual brains. This atlas (***Eickhoff et al., 2005***) allows assigning probabilistic values to in vivo brain images and has been used to, for example, validate the delineation of PAC on the basis of in vivo MRI images whose contrast is related to myelin (***Dick et al., 2012***).

The probabilistic atlas is generated using a volume registration method. Instead, the exceptionally reliable correspondence between micro- and macro-anatomy known to be present in many cortical areas (***Turner, 2013***) has inspired the use of registration methods that rely on cortical surfaces and macro-anatomical landmarks such as the major gyri and sulci (i.e. curvature based alignment rather than the whole volumetric data (***Fischl et al., 1999, 2008; Frost and Goebel, 2012; Goebel et al., 2006***)). Surface based alignment methods have been shown to improve the accuracy of inter-individual registration in micro-anatomically defined primary motor cortex (***Fischl, 2013***), the human middle temporal area (hMT) (***Frost and Goebel, 2013***), and to improve the registration of a cyto-architectonic atlas of the ventral visual system (***Rosenke et al., 2018; Fischl et al., 2008***).

Curvature based alignment is also routinely used in studies investigating the functional and anatomical properties of auditory cortical areas. However, Heschl’s Gyrus substantially varies in shape across individuals and across hemispheres, and slight changes in the primary auditory cortex location have been reported in subjects with a typical morphological variation of the Heschl’s Gyrus (***Heschl, 1878; Rademacher et al., 1993; Hackett et al., 2001; Marie et al., 2015***).

Given this variation in superior temporal plane macro-anatomy across individuals and shift of micro-anatomical areas with macro-anatomy, it is debatable if curvature based alignment improves the correspondence of micro-anatomically defined auditory areas. Accordingly, here we applied curvature based alignment (abbreviated as CBA), as well as a procedure tailored to the temporal lobe by incorporating anatomical priors (abbreviated as CBA+), and reconstructed cortical surfaces from the data of ***Morosan et al. (2001***) in order to investigate to what extent maximizing macro-anatomical inter-individual alignment improves the overlap of micro-anatomically defined auditory cortical areas. Our results address if the use of CBA or CBA+ is justified when considering the functional properties of auditory cortical areas. Thereby, our results not only test the validity of the use of CBA in previous studies, but also offer CBA+ to improve across participant alignment of the superior temporal plane as a tool to the auditory community. We showcase this approach by applying CBA+ to an in vivo dataset collected at 7 Tesla and projecting the improved cyto-architectonic atlas onto functional and anatomical group maps. In addition, in order to contribute to the ongoing debate on the in vivo localization of auditory cortical areas (***Moerel et al., 2014; Besle et al., 2018***), we align the cyto-architectonic atlas (and in vivo data) to a recent temporal lobe parcellation based on in vivo measurements (***Glasser et al., 2016***).

## Results

We obtained cyto-architectonically labeled temporal cortical areas and post mortem MR images of ten brains (volumetrically aligned (rigid body) to the Colin27 space) used in the JuBrain cyto-architectonic Atlas (***Amunts and Zilles, 2015***). The cyto-architectonically labeled areas were TE 1.0, TE 1.1, and TE 1.2 from ***Morosan et al. (2001***), TE 2.1 and TE 2.2 from ***Clarke and Morosan (2012***), TE 3 from ***Morosan et al. (2005***), and STS 1 and STS 2 from ***Zachlod et al. (2020***). In order to perform cortex based alignment, the white matter - gray matter boundary was segmented in all ten post mortem brains. To obtain such segmentation, we have used a combination of image filtering techniques and a histogram based segmentation approach (***Gulban et al., 2018b***), which reduced the amount of required manual corrections (see Methods section). Cortical surfaces were reconstructed to perform three different types of group alignment methods. These methods were rigid body (i.e. considering surface sampling [compared to volumentric alignment] and rigid body registration), curvature based alignment (CBA) and curvature based alignment with anatomical priors (CBA+; including the anterior Heschl’s Gyrus, the superior temporal gyrus, the superior temporal sulcus, and the middle temporal gyrus as anatomical priors). We additionally compared these surface approaches to the original volumetric alignment in the Colin27 space. We have validated the performance of these methods by comparing the overlap between cyto-architectonic areas across individuals. We subsequently used CBA+ to create superior temporal cortical group maps of in vivo MRI (at 7T) measurements and to align them to the probabilistic cyto-architectonic atlas.

### Comparison between alignment methods

***Figure 1*** rows 1 and 3 show the averaged curvature maps after alignment with each of the surface approaches we used (i.e., rigid only that linearly coregisters the surfaces, standard CBA, and CBA tailored to the temporal lobe [CBA+]). In the temporal lobe, the increased sharpness of the average curvature maps indicates the improved correspondence of the macro-anatomical features in CBA and CBA+ compared to the rigid only alignment. Especially in the right hemisphere (third row in figure ***Figure 1***), an improvement of CBA+ over standard CBA is noticeable at the level of the Heschl’s Gyrus (indicated by a red circle). The improvement in alignment of the macro-anatomical features in the temporal lobes (left and right) is also visible when considering the folded average meshes of the ten brains in the post mortem dataset (i.e. average folded meshes, ***Figure 1*** rows 2 and 4). In absence of large macro-anatomical differences across the individuals, improved alignment should increase the 3D complexity (e.g. gyri and sulci appearing very clearly distinguishable) of the average folded mesh. Cortical curvature-based alignment procedures, however, may be affected when individual cortical macro-anatomy strongly deviates from the average morphology. In the post mortem sample we analyzed, we observed macro-anatomical variations across hemispheres of two types. First, following the characterization described in ***Kim et al. (2000***); ***Da Costa et al. (2011***), the number of Heschl’s Gyri varied. In particular, we observed 1, 1.5 and 2 Heschl’s Gyri in [5, 4, and 1, respectively] right hemispheres and [6, 2, and 2, respectively] left hemispheres. Second, we observed the presence of three hemispheres (one right and two left ones) whose single Heschl’s Gyrus was continuous at the anterior part of the anterior temporal convolution, resulting in a split superior temporal gyrus (i.e. interrupted by an intermediate sulcus between the anterior and posterior part with respect to the location of the Heschl’s Gyrus - ***Figure 13*** lower right panel). This rare morphological pattern was first described in ***Heschl (1878***) and was reported to occur % 10 of all brains inspected by Richard L. Heschl (110 of 1087 brains). It was ∼18 times more likely to occur on the left hemisphere in comparison to right (also see ***Rademacher et al., 1993***, for another reference to Heschl’s work in English). As expected, the tailored alignment we developed here results in a more prominently defined Heschl’s Gyrus in the average mesh, resulting from the correct alignment of the anterior Heschl’s Gyrus across individual hemispheres. In the split superior temporal gyrus cases, we defined the gyrus as continuous (i.e. bridging the intermediate sulcus). While this definition did not compromise the alignment of the anterior Heschl’s Gyrus, the impact of the approach we followed in the alignment of regions in proximity to the intermediate sulcus would require a larger sample on which to evaluate alignment separately according to this macro-anatomical variation (i.e. aligning separately individuals with a split/continuous superior temporal gyrus).

**Figure 1.**
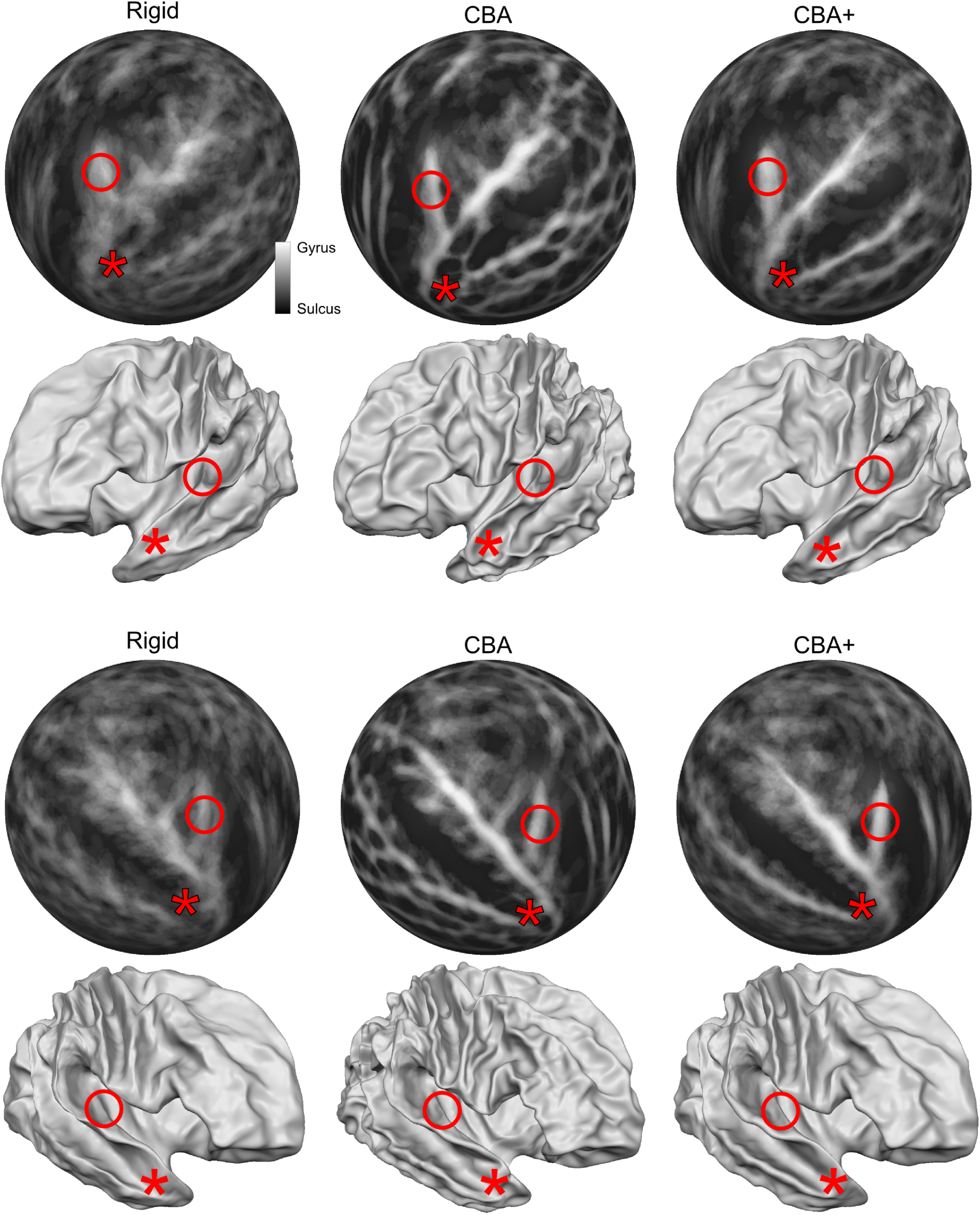
Differences between spherical rigid body alignment, curvature based alignment (CBA) and curvature based alignment with an anatomical prior (CBA+) on group average binarized curvature maps visualized as half-sphere projections (rows 1 and 3) and group average vertex coordinates visualized as folded surfaces (rows 2 and 4). In rows 1 and 3, higher contrast between sulci (dark gray) and gyri (light gray) shows more overlap around Heschl’s Gyrus which indicates that a method better accounts for inter-subject morphological variation. In rows 2 and 4, the average vertex coordinates show a more pronounced Heschl’s Gyrus in 3D as the alignment methods improves the anterior Heschl’s Gyrus overlap.

**Figure 2.**
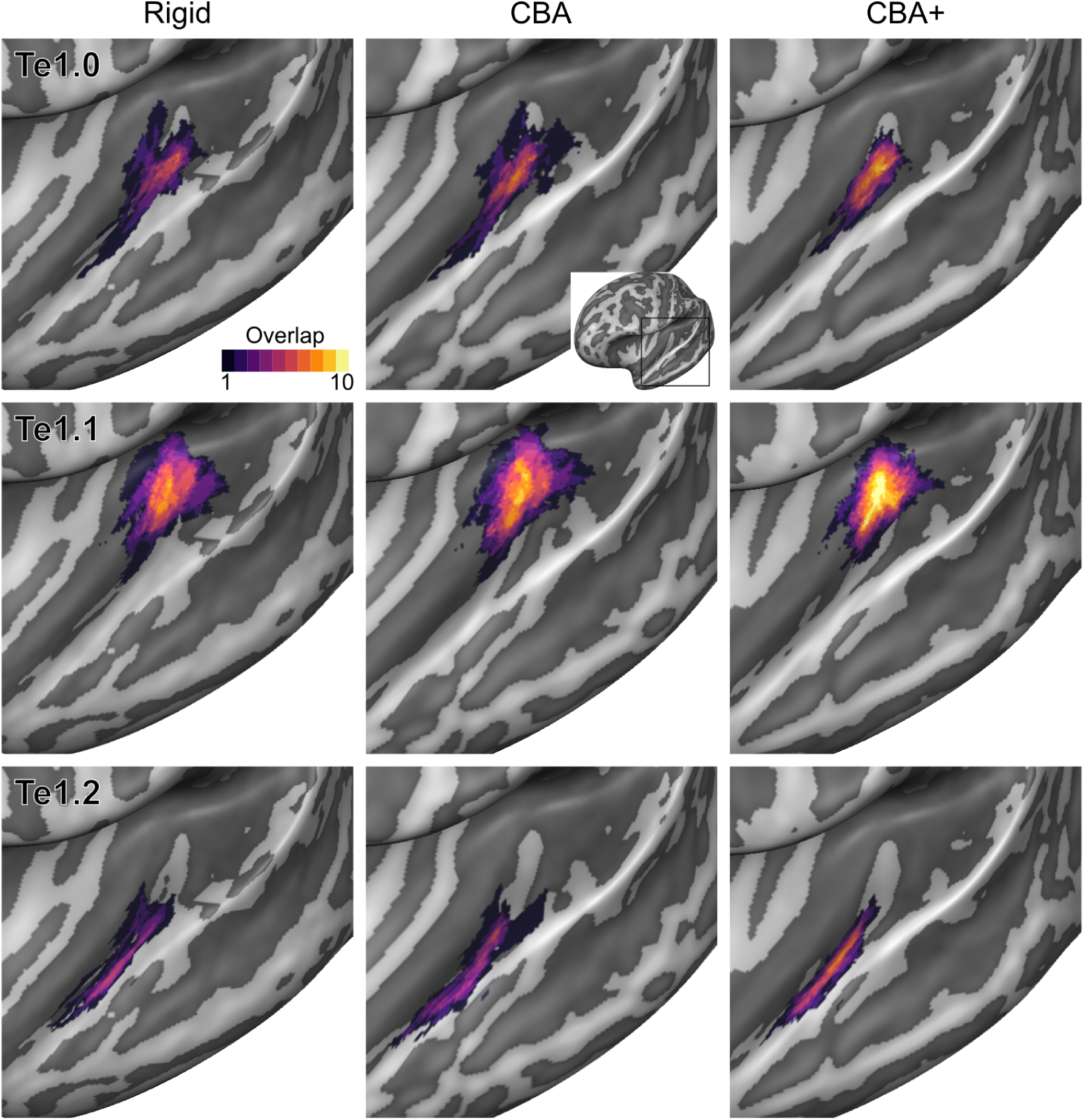
Probabilistic maps (after alignment) indicating the number of subjects for which a given vertex is labelled as belonging to the cyto-architectonic areas Te1.0, Te1.1 and Te1.2 are presented on inflated group average cortical surfaces of the left hemisphere. Columns show spherical rigid body alignment, curvature based alignment (CBA) and curvature based alignment with anatomical priors (CBA+) from left to right. Improvements in the micro-anatomical correspondence diminishes low values in the maps (purple) and increases the presence of high probability values (yellow).

**Figure 3.**
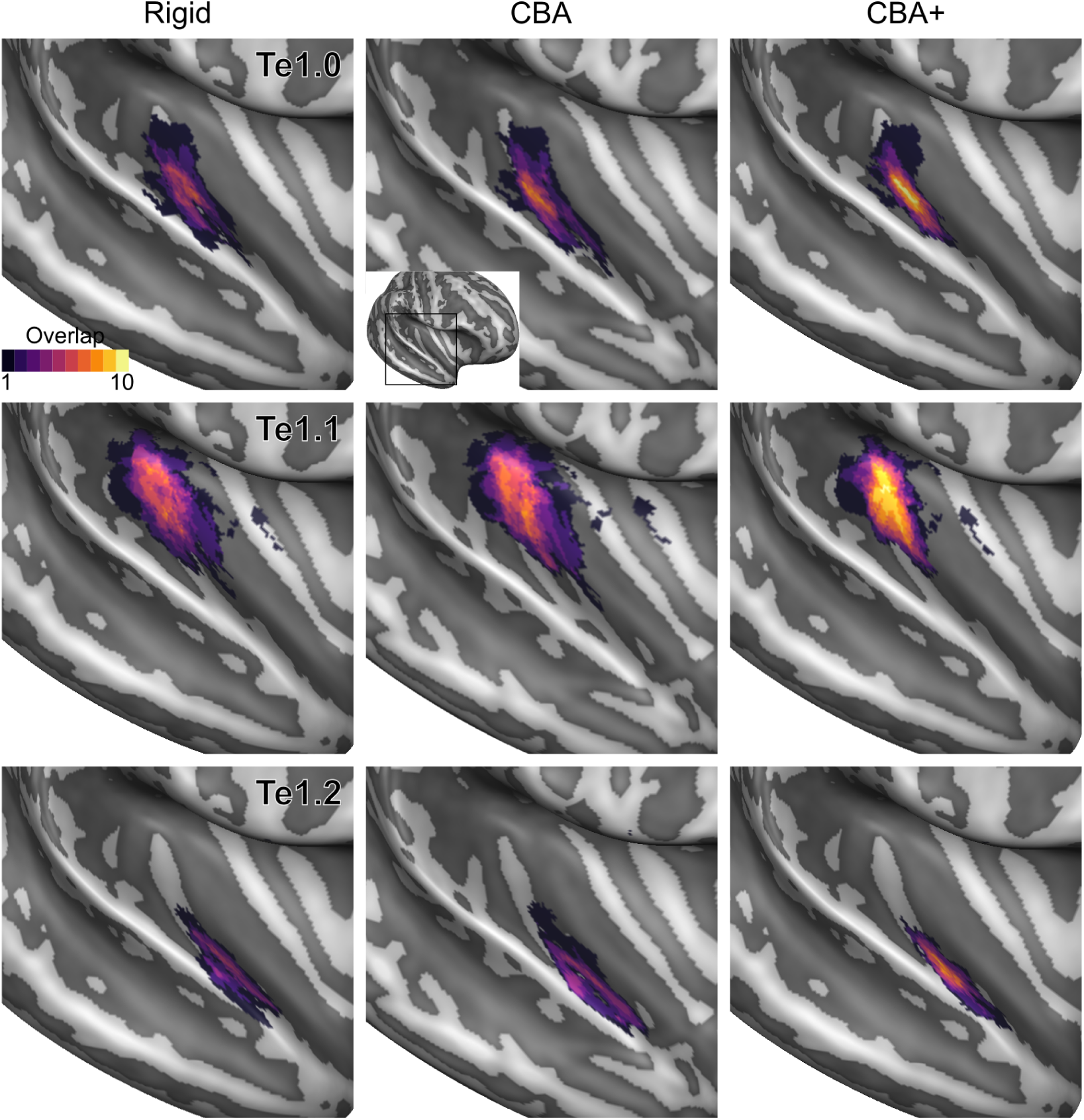
Probabilistic maps (after alignment) indicating the number of subjects for which a given vertex is labelled as belonging to the cyto-architectonic areas Te1.0, Te1.1 and Te1.2 are presented on inflated group average cortical surfaces of the right hemisphere. Columns show spherical rigid body alignment, curvature based alignment (CBA) and curvature based alignment with anatomical priors (CBA+) from left to right. Improvements in the micro-anatomical correspondence diminishes low values in the maps (purple) and increases the presence of high probability values (yellow).

**Figure 4.**
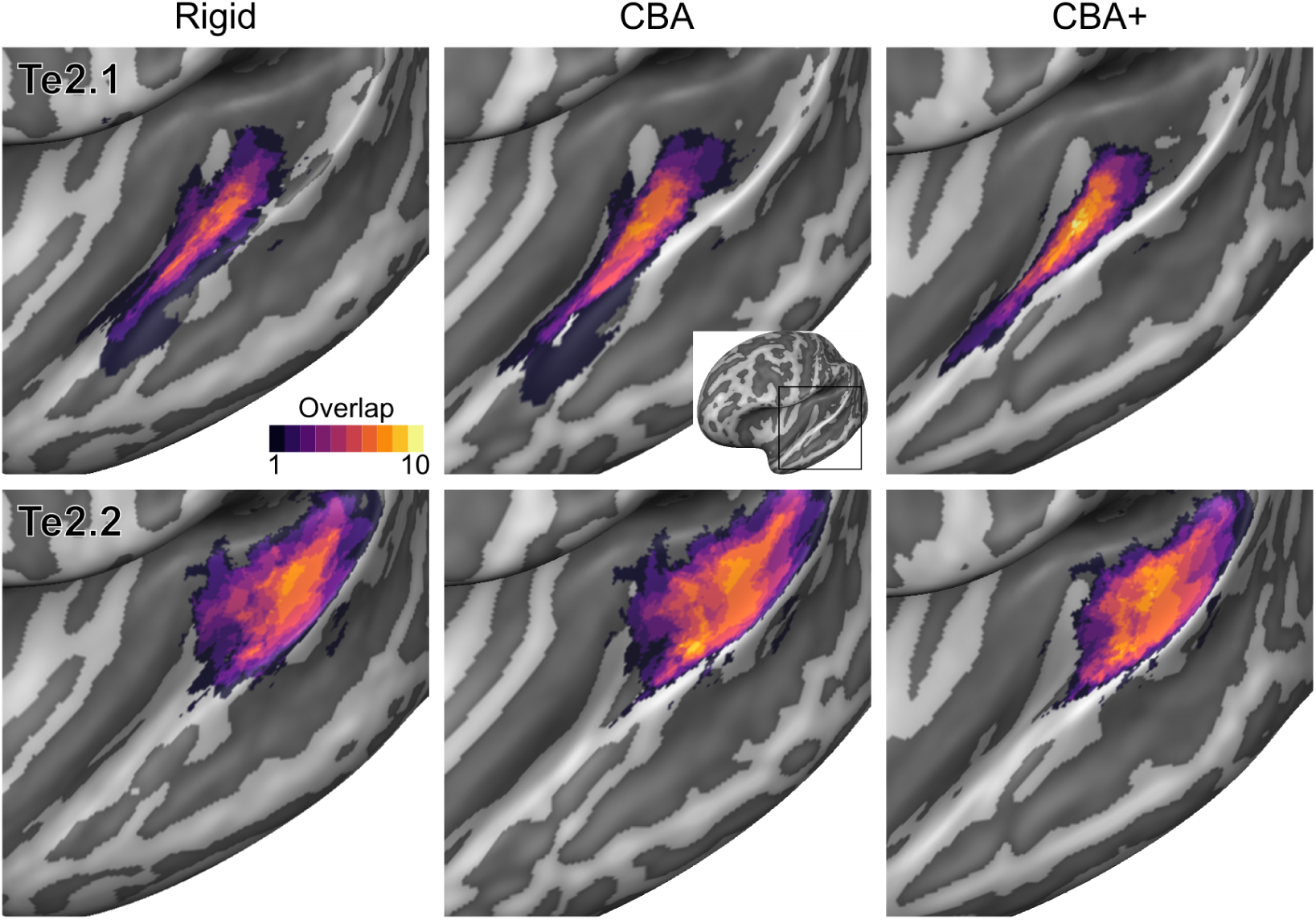
Probabilistic maps (after alignment) indicating the number of subjects for which a given vertex is labelled as belonging to the cyto-architectonic areas Te2.1 and Te2.2 are presented on inflated group average cortical surfaces of the left hemisphere. Columns show spherical rigid body alignment, curvature based alignment (CBA) and curvature based alignment with anatomical priors (CBA+) from left to right. Improvements the micro-anatomical correspondence diminishes low values in the maps (purple) and increases the presence of high probability values (yellow).

**Figure 5.**
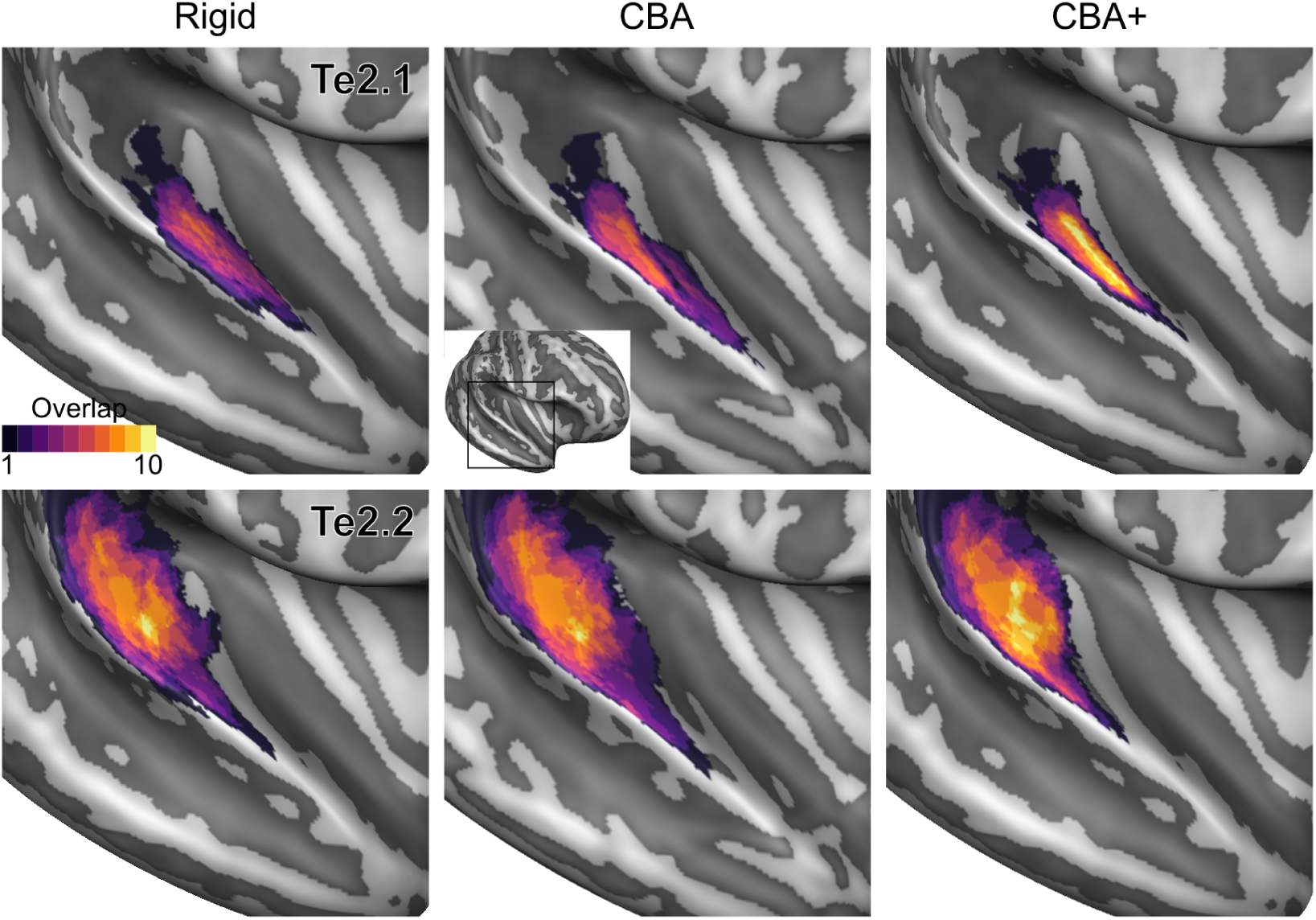
Probabilistic maps (after alignment) indicating the number of subjects for which a given vertex is labelled as belonging to the cyto-architectonic areas Te2.1 and Te2.2 are presented on inflated group average cortical surfaces of the right hemisphere. Columns show spherical rigid body alignment, curvature based alignment (CBA) and curvature based alignment with anatomical priors (CBA+) from left to right. Improvements in the micro-anatomical correspondence diminishes low values in the maps (purple) and increases the presence of high probability values (yellow).

**Figure 6.**
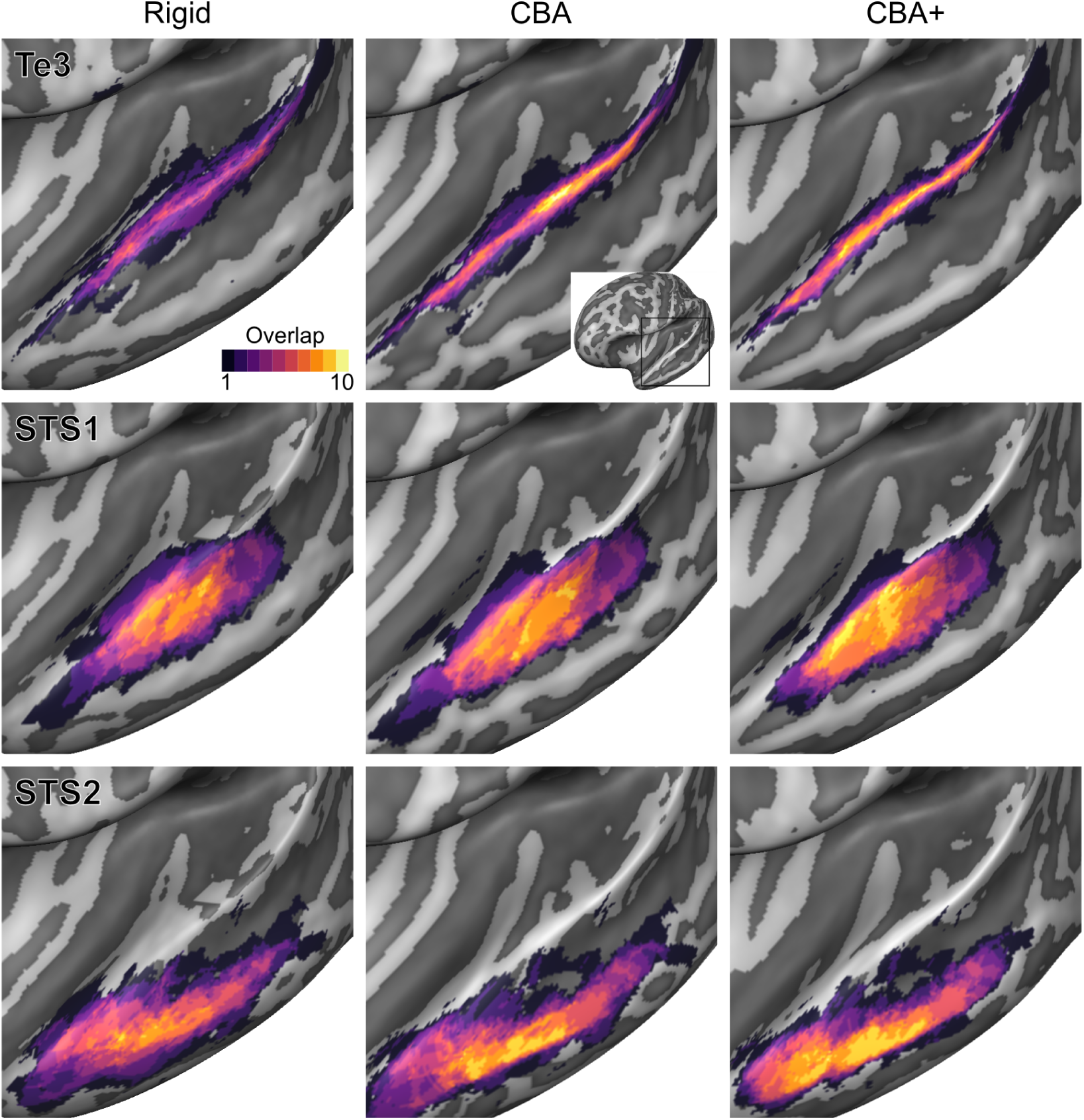
Probabilistic maps (after alignment) indicating the number of subjects for which a given vertex is labelled as belonging to the cyto-architectonic areas Te3, STS1 and STS2 are presented on inflated group average cortical surfaces of the left hemisphere. Columns show spherical rigid body alignment, curvature based alignment (CBA) and curvature based alignment with anatomical priors (CBA+) from left to right. Improvements in the micro-anatomical correspondence diminishes low values in the maps (purple) and increases the presence of high probability values (yellow).

**Figure 7.**
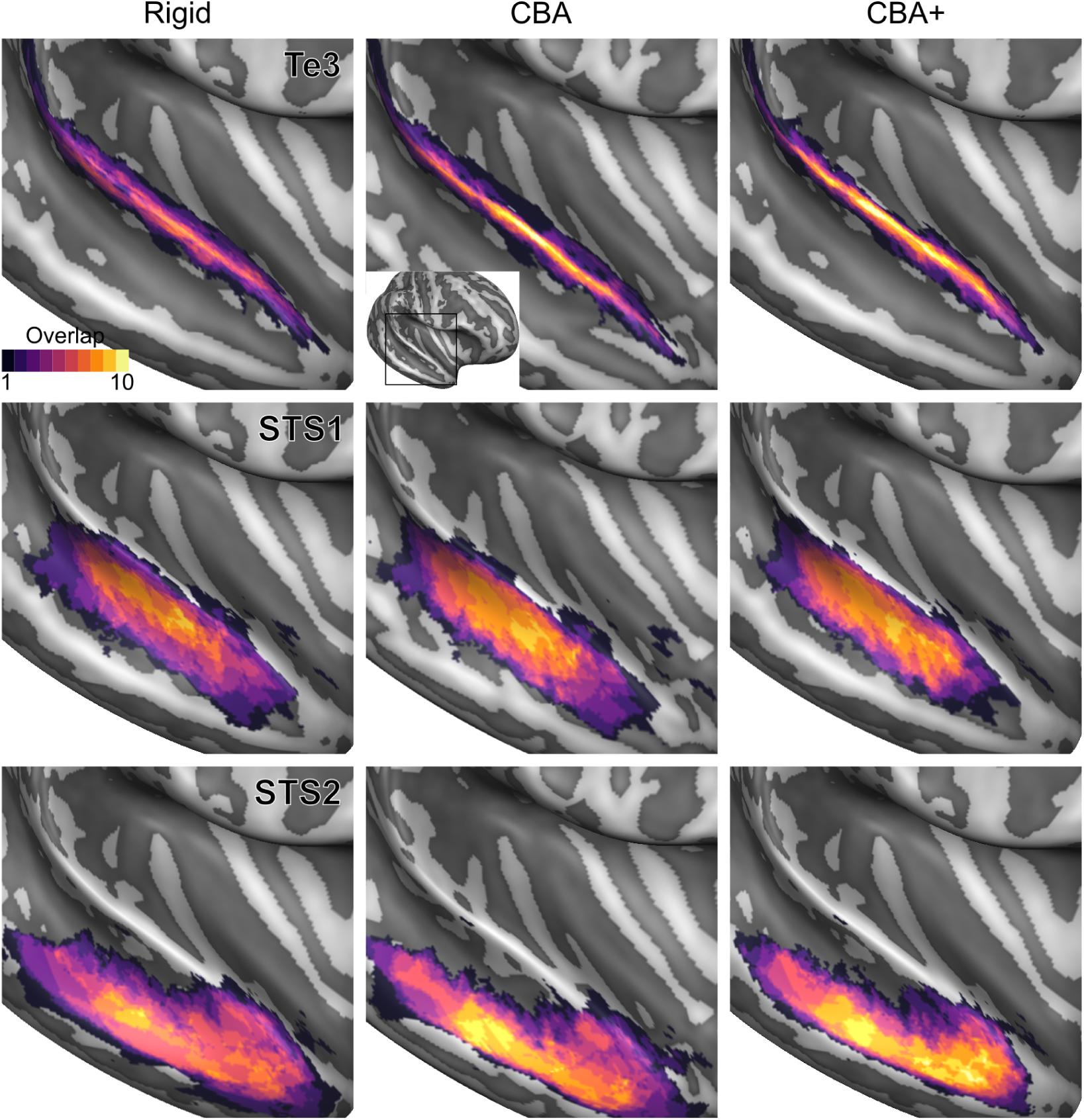
Probabilistic maps (after alignment) indicating the number of subjects for which a given vertex is labelled as belonging to the cyto-architectonic areas Te3, STS1 and STS2 are presented on inflated group average cortical surfaces of the right hemisphere. Columns show spherical rigid body alignment, curvature based alignment (CBA) and curvature based alignment with anatomical priors (CBA+) from left to right. Improvements in the micro-anatomical correspondence diminishes low values in the maps (purple) and increases the presence of high probability values (yellow).

**Figure 8.**
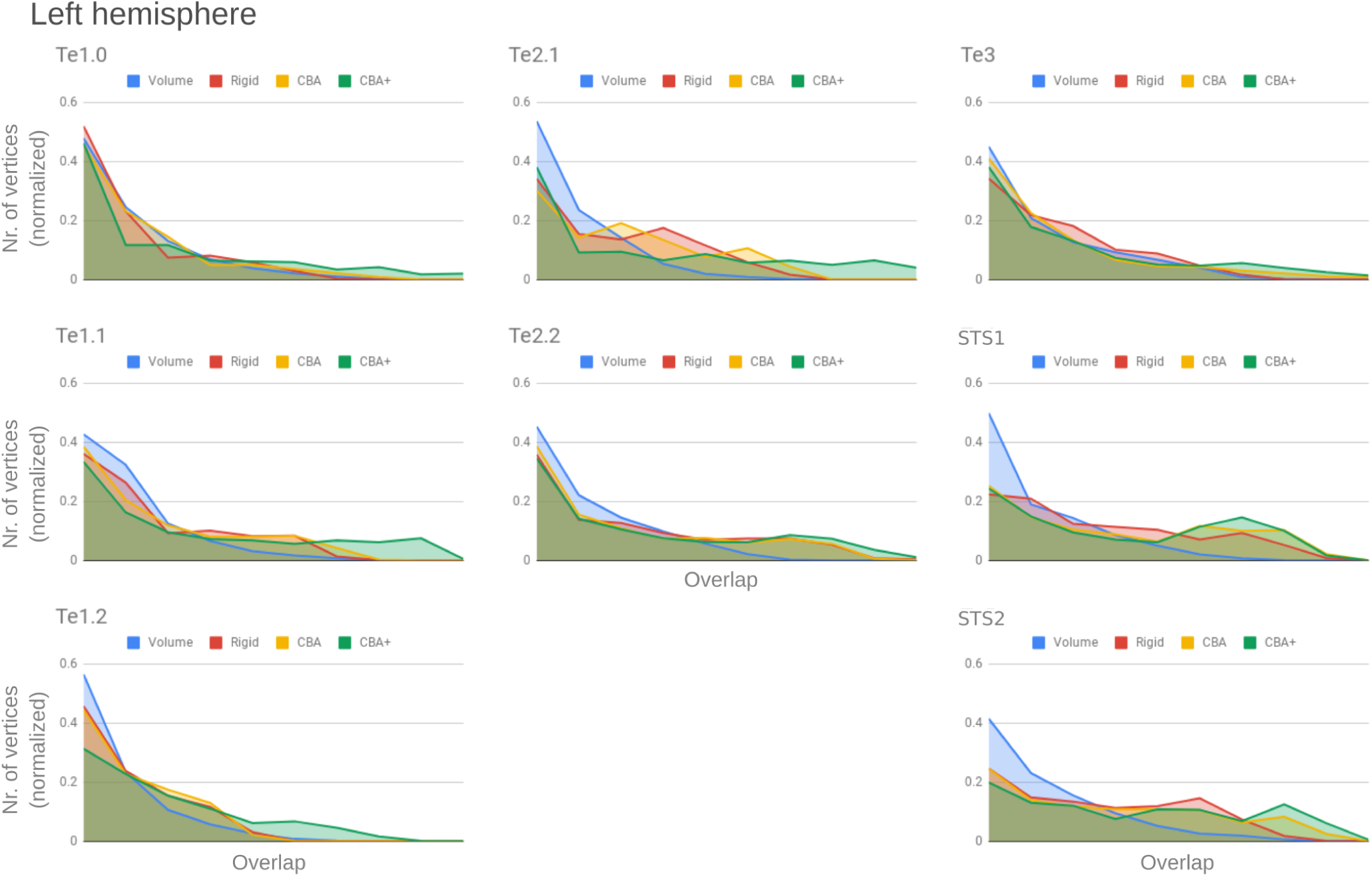
Histograms of the overlap across cyto-architectonic areas in the left hemisphere. The histograms are normalized by the number of vertices per area. The x-axis represents the probability value (an overlap from 1 out of ten [left] to 10 out of 10 participants [right]). The ideal co-registration method should show a less left skewed distribution. It can be seen that CBA+ shows the lowest skew towards the left in comparison to other methods.

**Figure 9.**
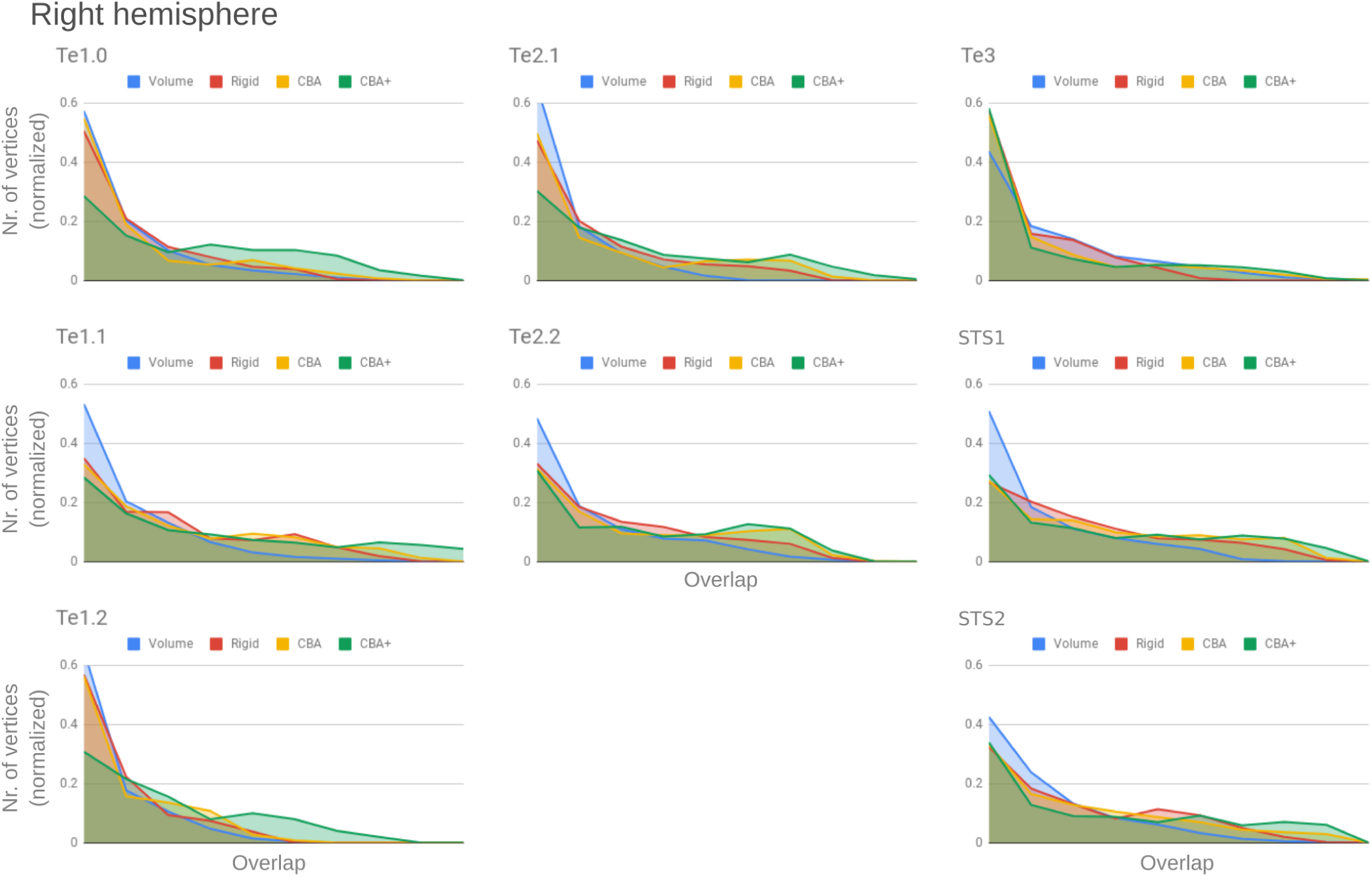
Histograms of the overlap across cyto-architectonic areas in the right hemisphere. The histograms are normalized by the number of vertices per area. The x-axis represents the probability value (an overlap from 1 out of ten [left] to 10 out of 10 participants [right]). The ideal co-registration method should show less left skewed distribution. It can be seen that CBA+ shows the lowest skew towards the left in comparison to other methods.

**Figure 10.**
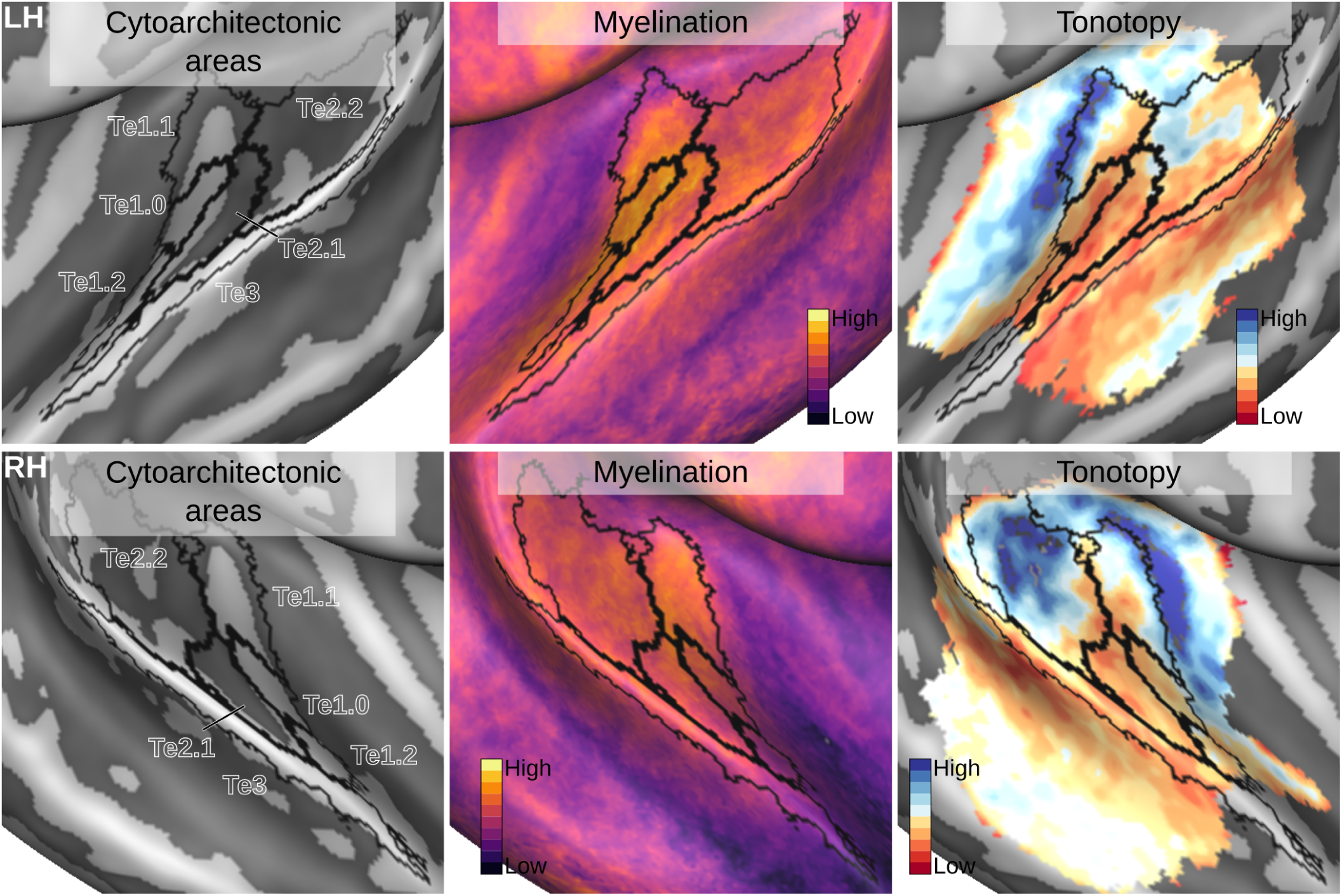
Relation between in vivo MRI measures and the cyto-architectonic atlas. The cyto-architectonic areas are delineated with black lines. The myelination index is computed from the division of T_1_w and 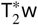 data. Tonotopy reflects the voxel-wise frequency preference estimated with fMRI encoding from the response to natural sound stimuli. All measures are sampled on the middle gray matter surfaces. **Figure 10–Figure supplement 1**. The same maps projected to an in vivo multi-modal MRI group atlas (***Glasser et al., 2016***).

**Figure 11.**
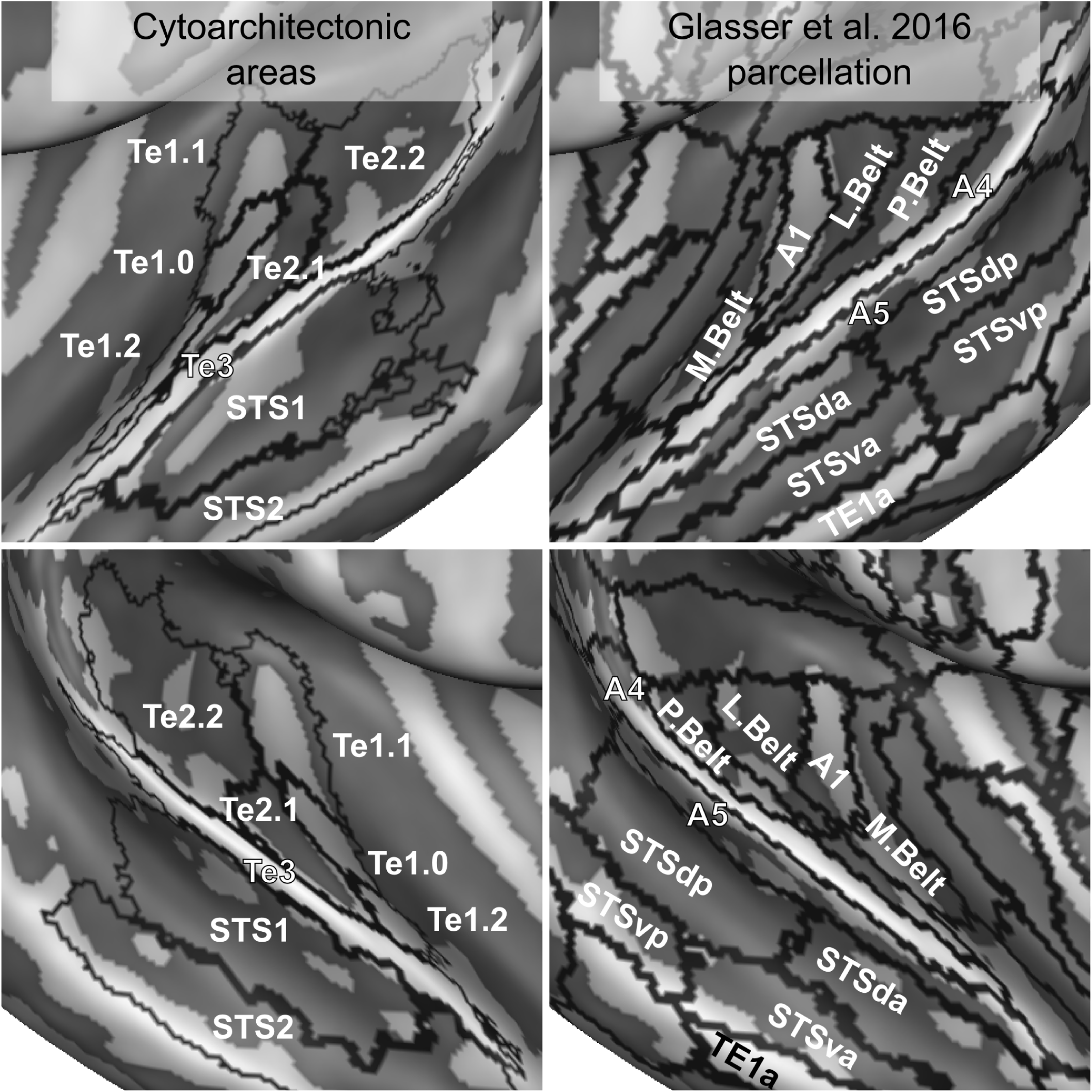
Comparison of cyto-architectonic areas ***Morosan et al. (2001***, 2005) and multi modal MRI based labels (***Glasser et al., 2016***). Areas on Heschl’s Gyrus differ between the two atlases.

**Figure 12.**
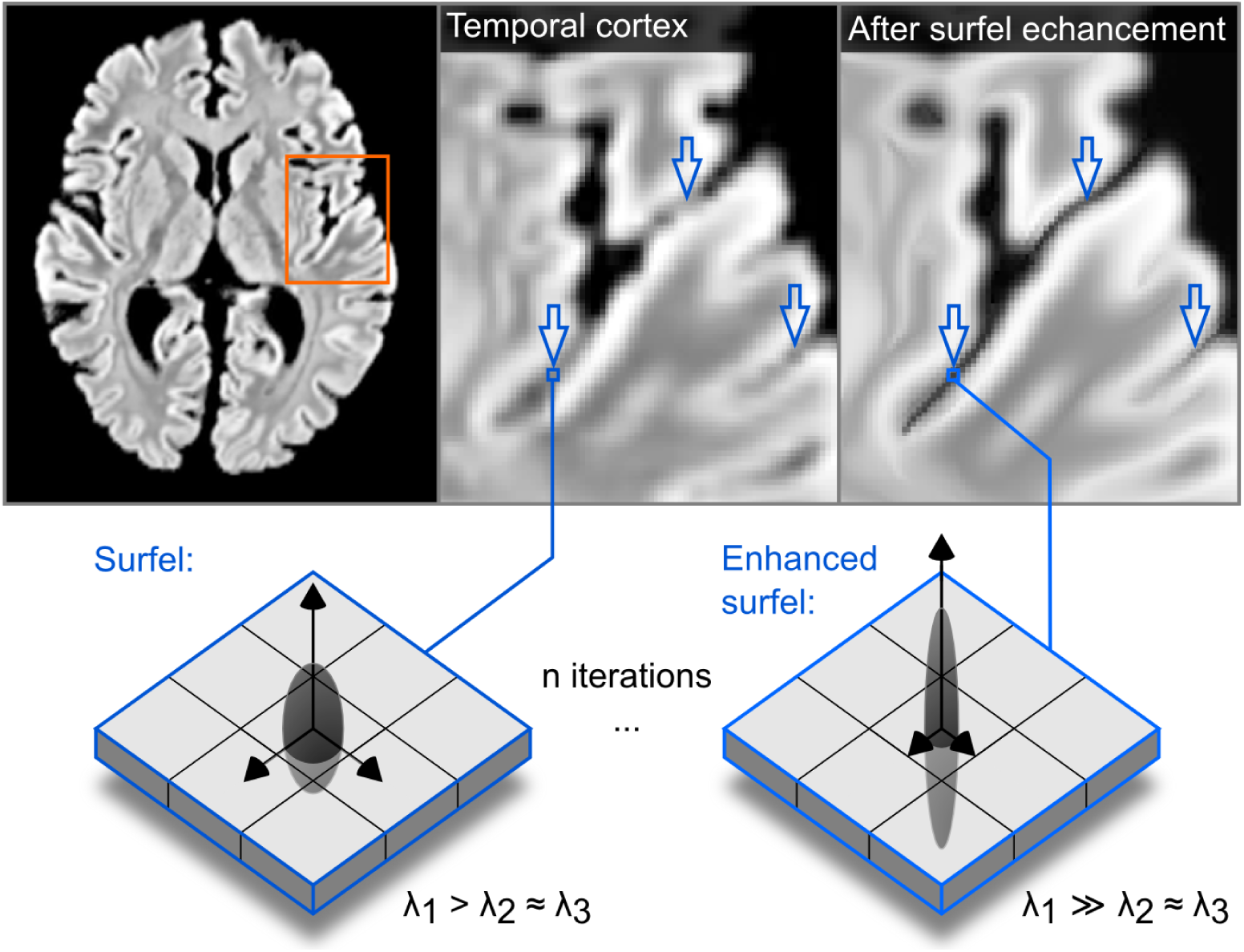
The effect of the structure enhancing filter shown on a transversal slice. Blue arrows point to locations where local contrast is sharpened.

**Figure 13.**
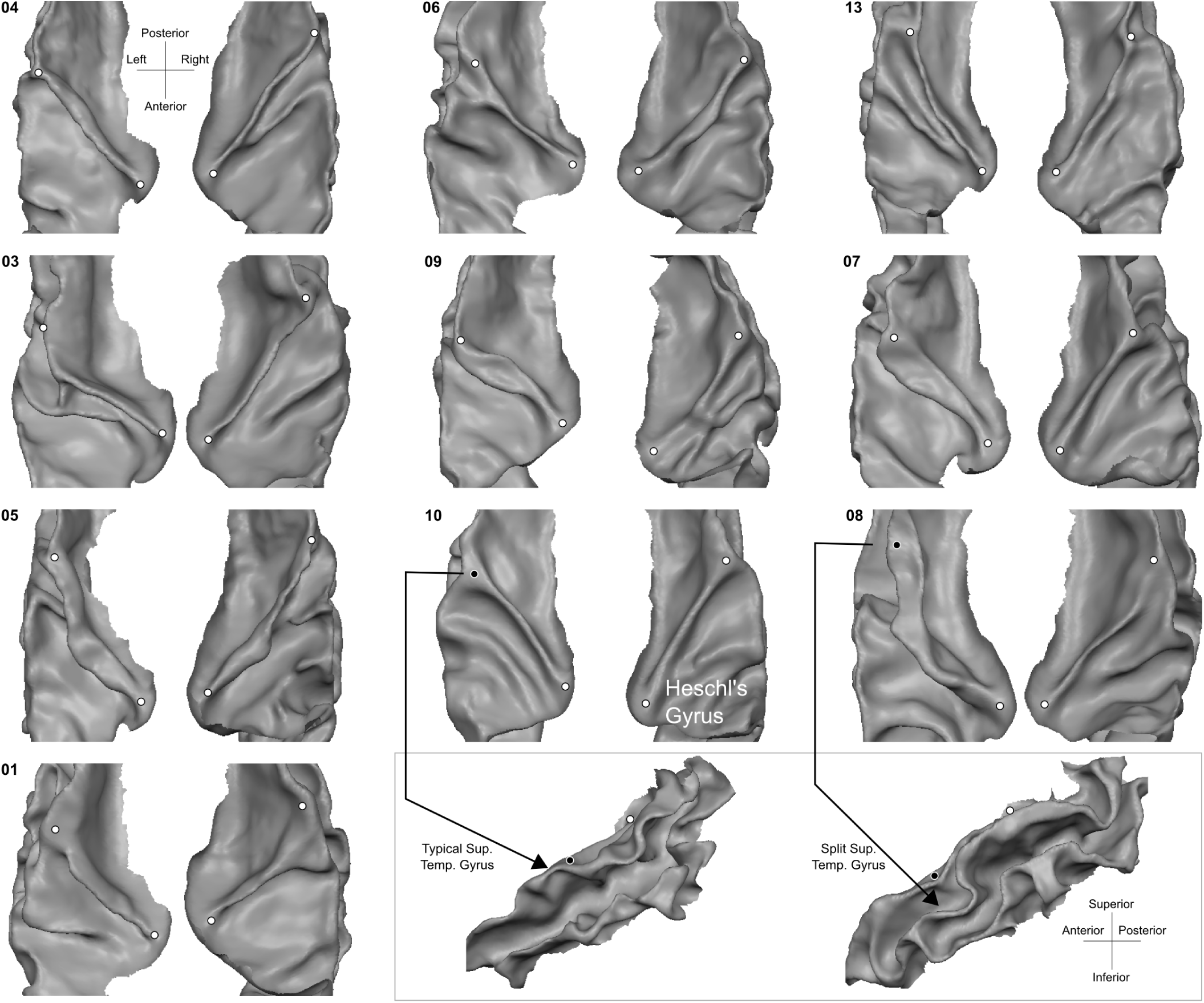
Individual superior temporal cortex white-gray matter boundary reconstructions. Anterior Heschl’s Gyrus is indicated as the gyrus between white dots. The bottom right side shows the rare occurrence of a split superior temporal gyrus (***Heschl, 1878***) in contrast to a typical superior temporal gyrus from the side view.

To evaluate the effect that minimizing macro-anatomical differences (as evidenced by the improved average curvature maps and folded meshes) has on micro-anatomy, we considered the inter-individual overlap of the cyto-architectonically-defined areas. In Figures 2-7 we present (for each labelled area) probabilistic maps (after alignment) indicating the number of subjects for which a given vertex is labelled as belonging to the same cyto-architectonic area. For all cyto-architectonic areas, CBA+ improves the overlap (as indicated by the increased probability of a vertex to be labelled as belonging to same area across the ten brains).

To better understand the differences between methods and quantitatively compare the rigid alignment, CBA, and CBA+ surface approaches to the initial volumetric alignment (in Colin27 space), ***Figure 8*** and ***Figure 9*** present the histograms of the probabilistic maps of each area (left and right hemisphere, respectively). For the cyto-architectonic areas along Heschl’s Gyrus (Te1.0, Te1.1 and Te1.2) the largest overlap is provided by CBA+, which improves micro-anatomical correspondence compared to the volume based alignment and the two other surface approaches we evaluated. For the areas in the planum temporale (Te2.1 and Te2.2), all surface approaches improve micro-anatomical correspondence compared to the volume alignment, and CBA+ provides an additional benefit especially for the area Te2.1. Similarly, for the areas in the superior temporal gyrus and sulcus and middle temporal gyrus (Te3, STS1 and STS2), all surface approaches improve micro-anatomical correspondence compared to the volume alignment while differences between standard CBA and CBA+ are modest.

### Aligning in vivo group measures to the probabilistic post mortem areas

The definition of probabilistic cyto-architectonically defined areas has been previously used to analyze in vivo functional and anatomical data (see e.g. (***Dick et al., 2012***)). Here we demonstrate the use of CBA+ and the improved version of the cyto-architectonic atlas to this end. In particular, we aligned in vivo data collected at 7 Tesla to the CBA+ aligned post mortem cyto-architectonic atlas. We considered only the areas in the superior temporal cortex (Te1.0, Te1.1, Te1.2, Te2.1, Te2.2 and Te3) as they were consistently included in the imaged field of view in the in vivo dataset. First, we used CBA+ to produce an average morphology for the in vivo data. This alignment allowed us to derive group level maps based on the available anatomical and functional data. In particular, anatomical MRI data (0.7 mm isotropic) were used to derive intra cortical contrast related to myelin from the division of T_1_w and 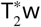 data. In addition, functional MRI data (1.1 mm isotropic) collected by presenting natural sounds and analyzed with an fMRI encoding approach (***Moerel et al., 2012***), were used to derive tonotopic maps (see ***Figure 10***. Second, using CBA+, we aligned the average morphology of the in vivo data to the cyto-architectonic atlas. This allowed us to project cyto-architectonic parcels on the in vivo maps and evaluate their relationship.

Intra-cortical contrast related to myelin highlights the (medial) Heschl’s Gyrus as the most myelinated region in the temporal cortex (see ***Figure 10***). Across cyto-architectonic areas, Te1.0 shows the highest myelination contrast. Myelin related contrast is also high in the most medial portion of Heschl’s Gyrus (Te1.1) and gradually decreases when moving away from Heschl’s Gyrus.

The average tonotopic pattern highlights the Heschl’s Gyrus as, for the most part, preferring low frequencies, while surrounding areas (in posterior antero-medial and antero-lateral directions) prefer high frequencies (see ***Figure 10***). The high frequency areas form an inverted “V” pattern surrounding the Heschl’s Gyrus (***Da Costa et al., 2011; Moerel et al., 2014***). Cyto-architectonic primary cortical areas (Te1) cover the Heschl’ gyrus, with the core (Te1.0) in its middle section which (at the group level) appears characterized by mainly low frequency preference (see ***Figure 10***). Located medial to Te1.0, area Te1.1 may reflect an intermediate processing stage between primary and belt areas (***Moerel et al., 2014***) and covers one tonotopic gradient going from high to low on an antero-medial to postero-lateral direction. Te2.2 covers a posterior portion of the tonotopic gradient running in the posterior to anterior direction. Te2.1, covering an intermediate location between Te2.2 and Te1.0/Te1.2, overlaps with a low frequency preferring region in the lateral portion of the Heschl’s sulcus. Finally, Te3 covers a low frequency portion of the tonotopic maps along the superior temporal gyrus (***Moerel et al., 2014***). For comparison, in a supplement to ***Figure 10*** we report the same maps aligned with an an atlas obtained from in vivo MRI data (using both anatomical and functional information) in a large cohort (***Glasser et al., 2016***). A direct comparison between the post mortem and in vivo atlases projected on the average anatomical curvature of our in vivo data is reported in ***Figure 11***.

## Discussion

The superior temporal plane shows considerable macro-anatomical variability across individuals (***Pfeifer, 1921, 1936; Von Economo and Horn, 1930; Rademacher et al., 1993; Zoellner et al., 2019***). Here we evaluated the effect of macro-anatomical variability on localizing cyto-architectonic areas across different brains. We have used ten individual brains available from the JuBrain cyto-architectonic Atlas^1^ (***Amunts and Zilles, 2015***) together with a surface registration method that minimizes macro-anatomical variability around the transverse temporal gyrus (similar to ***Rosenke et al., 2018***) to show that minimizing macro-anatomical variability in the superior temporal plane results in improved micro-anatomical correspondence across brains.

Applying a surface registration for inter-subject alignment required accurate segmentation of the post mortem MRI dataset. While this issue has been tackled before for the investigation of cyto-architectonic areas in the visual cortex ***Rosenke et al. (2018***), an accurate segmentation of the temporal areas was not available. To obtain such segmentation and reduce the amount of manual corrections, we have used a tailored procedure based on image filtering and histogram based segmentation (***Gulban et al., 2018b***). The resulting segmentations allowed us to define the macro-anatomical variability in the sample (see ***Figure 13***). The availabe ten brains showed typical variations in the morphology of the Heschl’s Gyrus (with a single Heschl’s Gyrus being the most prevalent one), as well as cases in which the Heschl’s Gyrus was continuous to the anterior portion of the superior temporal gyrus (***Heschl, 1878***).

The segmented hemispheres were used for cortex based alignment. The standard approach minimizes macro-anatomical variation across subjects (i.e. maximizes the overlap of the curvature maps) across the whole brain (in a coarse to fine iterative approach). As such, standard CBA is driven by the major anatomical landmarks including the superior temporal gyrus and sulcus. This, however, can result in compromised alignment of smaller (but consistent) anatomical features such as the Heschl’s Gyrus. This can be seen in ***Figure 1*** (middle column) where the compromised alignment of the Heschl’s Gyrus across hemispheres is indicated by the reduced sharpness of the averaged binarized curvature maps. For this reason, here we have considered the application of an approach tailored to the superior temporal plane. By providing additional landmarks (the Heschl’s Gyrus, the superior temporal gyrus/sulcus and middle temporal gyrus) to the CBA procedure, we improved the alignment across subjects in the superior temporal cortex (see e.g. the difference in the average curvature maps between standard CBA and CBA+ in ***Figure 1***). Both the CBA and CBA+ approach greatly improved the macro-anatomical correspondence when compared to a rigid body procedure (which by sampling the volumetric data on surfaces already offers an improvement compared to the original volumetric alignment - see ***Figure 8*** and ***Figure 9***). The advantage for the tailored approach (CBA+, rightmost column in ***Figure 1***) is stronger on the right hemisphere, with some residual misalignment for the left Heschl’s Gyrus. This difference in performance could be explained by the larger prevalence (within our sample) in the left hemisphere of cases with the Heschl’s Gyrus merging with the anterior portion of the superior temporal gyrus (i.e. split superior temporal gyrus cases; two in the left and one in the right hemisphere). In the future, a larger sample could allow evaluating this issue, as well as the impact that the inclusion of this macro-anatomical variation has on the alignment of regions close to the superior temporal gyrus, by evaluating the alignment separately (with and without) such cases.

Improving macro-anatomical correspondence resulted in improved overlap of the cyto-architectonic areas across subjects. As a result of the CBA+ alignment, the micro-anatomically defined areas were smaller and the probability for a vertex to be labelled as belonging to the same area across the post mortem samples was higher (see Figures 2-7 and the histograms in ***Figure 8*** and ***Figure 9***). The tailored approach (CBA+) resulted in increased overlap (also compared to standard CBA) in all areas but especially for those on Heschl’s Gyrus or immediately adjacent to it (Te1.0, Te1.1, Te1.2 and Te2.1). This result is a direct consequence of defining the (most anterior) Heschl’s Gyrus as an additional landmark for alignment. The most anterior Heschl’s Gyrus was recognized as the putative location of primary auditory cortex in the case of complete duplication on the basis of myelo-architecture (***Hackett et al., 2001***). When this anatomical landmark is not used, the duplication of the Heschl’s gyrus results in poorer matching across subjects (i.e., the most posterior duplication of some subjects is aligned to the single Heschl’s Gyrus of other subjects). The post mortem dataset includes six Heschl’s Gyrus duplication cases (four in the right and two in the left hemisphere). Follow up studies are needed to evaluate the effect of an incomplete duplication of Heschl’s Gyrus. As previous myelo-architecture studies reported a shift of primary areas towards the intermediate Heschl’s sulcus in the case of an incomplete duplication (***Hackett et al., 2001***), a partial alignment of the primary areas (Te1.0 and Te1.1) may be expected. Examining the effect of an incomplete duplication on micro-anatomical alignment provide additional insights for a further refinement of the alignment procedure we propose here. In addition to the anterior Heschl’s Gyrus, CBA+ includes the superior temporal gyrus/sulcus and middle temporal gyrus as anatomical landmarks. While to a lesser degree than the areas on Heschl’s Gyrus, areas along these landmarks were also better realigned by CBA+. This indicates that favoring these gyri/sulci with respect to other major landmarks on the cortex is beneficial for the alignment of temporal areas. The improved cyto-architectonic overlap obtained with CBA+ suggests that this approach may be relevant for the functional and anatomical investigation of (auditory) temporal areas in vivo, as well as the investigation (post mortem and in vivo) of other cortical regions in which macro anatomical variability is high. We make the individual hemisphere surface models and the individual cyto-architectonic areas used in this study publicly available at https://kg.ebrains.eu/search/instances/Dataset/ff71a4d1-ea14-4ed6-898e-b92d95b3c446.

To showcase the application of CBA+ to the analysis of in vivo MRI data, we applied the same procedure to align anatomical and functional data collected at 7 Tesla across individuals. In addition, we used CBA+ to align the in vivo data to the improved cyto-architectonic atlas.

The pattern of myelin related intra-cortical contrast followed previous reports (***Glasser and Van Essen, 2011; Dick et al., 2012; De Martino et al., 2015***). The alignment to the cyto-architectonic atlas shows a high myelin related contrast in area Te1.0, in agreement with previous studies (***Dick et al., 2012***). Myelin related contrast was high also in the most medial portion of Heschl’s Gyrus (Te1.1) and decreased when moving away from Heschl’s Gyrus. While subtle differences between Te1.0 and Te1.1 were already noticeable, a more clear cut separation between these regions may require the evaluation of myelin related contrast across depths similarly to previous approaches (***Dick et al., 2012; De Martino et al., 2015***). In addition, future investigations may evaluate the information provided by intra anatomical contrast resulting from in vivo MRI acquisitions other than the one we considered here. For instance using the orientation of intra cortical fibres (***McNab et al., 2013; Gulban et al., 2018a***).

The group tonotopy maps we derived from the in vivo data follow previous reports (***Dick et al., 2012; Moerel et al., 2014; Besle et al., 2018***). In particular, they show one gradient within area Te1.1 progressing from high to low frequencies in antero-medial to postero-lateral direction. Based on the average maps, a full tonotopic gradient was not visible in Te1.0, which was corresponding mainly with the low frequency area in medial Heschl’s Gyrus. This pattern may be the result of excessive smoothing caused by inter-subject averaging that highlights the larger frequency gradient that in tonotopic maps progresses in the anterior-posterior direction on the planum temporale and thus favors the interpretation of the pattern within larger cortical areas (***Moerel et al., 2014***). More fine grained information (within smaller areas such as e.g. Te1.0) could be leveraged by considering single subjects in the future (***Moerel et al., 2014***). Te2.2 captured the most posterior portion of the larger tonotopic gradient that, consistently with previous reports, we identify as running in a direction orthogonal to Heschl’s Gyrus (***Moerel et al., 2014; Besle et al., 2018***). The other cyto-architectonic regions that overlapped with our functional acquisition field of view (Te2.1 and Te3) covered low frequency preferring regions of the tonotopic map in the lateral portion of the Heschl’s sulcus and the superior temporal gyrus. These results argue for the necessity of interpreting large scale tonotopic maps which alone do not allow defining the borders between cortical areas (***Moerel et al., 2014***). A large tonotopic gradient unarguably runs in a posterior to anterior direction ***Da Costa et al. (2011***); ***Besle et al. (2018***). Equating this gradient with the gradient that identifies the primary auditory cortex results in a view in which the core lies orthogonal to Heschl’s Gyrus (***Da Costa et al., 2011; Saenz and Langers, 2014; Besle et al., 2018***). On the other hand, the cyto-architectonic areas-now restricted in size by better aligning macro-anatomy-suggest that the auditory core (Te1) runs along Heschl’s Gyrus (i.e. the “classical” view; (***Dick et al., 2012; Moerel et al., 2014***)). This view is strengthened by the combined interpretation of myelin and tonotopy (see ***Figure 10*** and results in (***Dick et al., 2012; Moerel et al., 2014***)) as well as other auditory cortical functional characteristics (e.g., frequency selectivity; (***Moerel et al., 2014***)).

Interesting differences exist between the surface projection of the cyto-architectonic areas compared to a recent parcellation of the temporal lobe derived solely from in vivo imaging (***Glasser et al. (2016)*** - see ***Figure 11***). Cyto-architectonic areas Te1.1, Te1.0 and Te1.2 lie postero-medial to antero-lateral along the Heschl’s Gyrus. The most lateral subdivision (Te1.2) has been suggested to be the human homologue of area RT in the monkey (and thus part of the auditory core) or part of the lateral belt (***Moerel et al., 2014***). In the multi modal MRI parcellation, on the other hand, Heschl’s Gyrus is divided in an area labelled as A1, corresponding to the most medial two thirds, and its most lateral portion, which is part of the area labelled as the medial belt. Outside of the Heschl’s Gyrus other differences between the in vivo and post mortem atlas are visible. The lateral belt and parabelt areas as defined in the in vivo atlas occupy an area roughly corresponding to Te2, but the border between the areas labelled as belt and parabelt run approximately orthogonal to the border between Te2.1 and Te2.2. Te3, previously considered as an homologue of parabelt, corresponds to the areas labelled as A4 and A5 in the in vivo atlas. STS1 overlaps with the dorsal portion of superior temporal sulcus (STSda and STSdp in the in vivo atlas) and STS2 with the ventral portion of superior temporal sulcus for the most part. While the in vivo multi modal atlas has been derived from a large sample of participants (N=210), these differences may be caused by an insufficient amount of information available in the in vivo data used for the parcellation of the superior temporal plane.

In conclusion, here we show that an alignment procedure tailored to the superior temporal cortex and driven by anatomical priors together with curvature values improves inter-subject correspondence of cyto-architectonic areas. Reducing macro-anatomical variability and improving cyto-architectural correspondence may reduce the inter-subject variability of (anatomical and functional) characteristics probed in vivo, resulting in a more accurate definition of putative cortical (temporal) areas. Thereby our tailored approach has the potential to improve the investigation of anatomical and functional characteristics of auditory cortical areas using in vivo MRI. While we demonstrate its effectiveness in the temporal cortex, this approach is easily extendable to other cortical areas in which macro-anatomical inter subject variability is not easily accounted for by standard surface registration methods. Future studies should evaluate if this procedure, apart from being more accurate, is equally accurate for all known macro-anatomical variations of the morphology of the Heschl’s Gyrus. To demonstrate its applicability in vivo, we used CBA+ on data collected at 7 Tesla and coregistered our data to the post mortem atlas. In future work, CBA+ may aid the parcellation of the auditory cortex based as well as the other brain regions (e.g. frontal cortex) on in vivo data.

## Methods

### Post mortem data

We used the cyto-architectonically labeled temporal cortical areas of the ten brains used in ***Morosan et al. (2001***, 2005); ***Zachlod et al. (2020***). The labeled areas were Te1.0, Te1.1, Te1.2, Te2.1, Te2.2 (***Morosan et al., 2001***), Te3 (***Morosan et al., 2005***), STS1, STS2 (***Zachlod et al., 2020***). All brains were linearly registered to Colin27 space (***Evans et al., 2012***) at 1 mm isotropic resolution, which was the starting point for all further analyses.

#### Cortical segmentation

In order to perform cortex based alignment, the white matter - gray matter boundary was segmented in all ten post mortem brains. The anatomical image quality was insufficient to employ fully automatic segmentation methods. To mitigate this issue, we employed a spatial filter that was applied to an upsampled version of the data (to 0.5mm isotropic). This spatial filter was tailored to exploit the structure tensor field derived from the images. Our implementation of this procedure-that mostly follows ***Weickert (1998***); ***Mirebeau et al. (2015***)-included the following steps:

1. Smoothing the image for spatial regularization

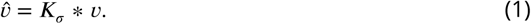

where * indicates convolution and *K* is a Gaussian kernel with standard deviation defined by *σ*. Here we have opted for *σ* = 1.
2. Computing the gradients of the image to obtain a vector field (we have used central differences)

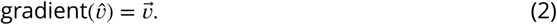
3. Generating a structure tensor field by using the self outer product:

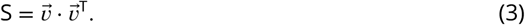
4. Decomposing (using eigen decomposition) the structure tensor field:

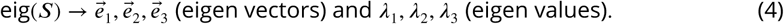 Note that the eigen vectors are sorted according to eigen values *λ*_1_ > *λ*_2_ > *λ*_3_.
5. Using eigen values to derive a vector field:

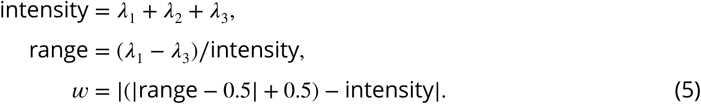 Here we wanted to enhance prolate ellipsoid tensors (also called surfels, surface elements, *λ*_1_ ≈ *λ*_2_ > *λ*_3_) more than isotropic structure tensors (*λ*_1_ ≈ *λ*_2_ ≈ *λ*_3_) and oblate ellipsoid tensor (also called curvels, curve elements like tubes, *λ*_1_ > *λ*_2_ ≈ *λ*_3_).
6. Generating a diffusion tensor field from weighted eigen vectors:

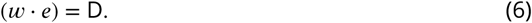
7. Smoothing the diffusion tensor field.

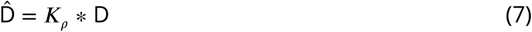

where * indicates convolution and *K* is a 3D Gaussian kernel with *σ* standard deviation. Here we have used *ρ* = 1. A higher value would enhance features at a larger spatial scale.
8. Computing a vector field (the flux field) using the diffusion tensor field and eigen vectors (D_*i*_ is a tensor 3× 3; 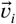 is a vector 1× 3):

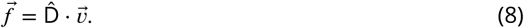
9. Updating the image (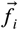 is a vector; *v*_*i*_ isa scalar):

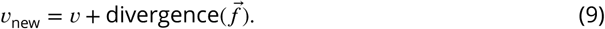
10. Repeating all steps until the desired number of iterations is reached (each iteration diffuses the image more and the diffusion is non-linear and anisotropic).

For segmenting the post mortem data, here we iterated this process 40 times. This number of iterations was visually judged as sufficient to enhance the boundary between white matter and gray matter as well as distinguishing the two banks of sulci by rendering them sharper [see ***Figure 12***]. Our implementation is available within the Segmentator package version 1.5.3 (***Gulban and Schneider, 2019***).

#### Cortical surface reconstruction

After filtering the images, we generated an initial white matter segmentation using intensity-gradient magnitude joint 2D histograms (***Gulban et al., 2018b***). This initial segmentation was corrected in two stages. First, manual corrections were performed by O.F.G using both enhanced and un-enhanced anatomical images (around 8 hours of manual work per brain). Second, after splitting left and right hemispheres, we generated surfaces as triangular meshes using the marching cubes method (as implemented in BrainVoyager 21.4, ***Goebel (2012***)) and decimating the total amount of vertices to 200000 (with approximately equal edge lengths). The surfaces were visually checked for bridges and holes and problematic areas were corrected until the Euler characteristic of each surface became 2 (i.e. topologically identical to a sphere). ***Figure 13*** shows the morphological variation across the post mortem brains on the superior temporal cortex.

#### Cortical surface alignment

The prepared surfaces were inflated to an approximate sphere and mapped onto a high density spherical mesh (163842 vertices). Prior to cortex based alignment, the meshes were aligned using a spherical rigid body method to minimize curvature differences across subjects [see ***Figure 1*** left column]. Cortex based alignment was performed in two different ways. First, we non linearly registered the surfaces of each hemisphere across brains using standard cortex based alignment (i.e. minimizing curvature differences across individuals in a coarse to fine manner (***Frost and Goebel, 2012***)). Second, to tailor the alignment to the superior temporal cortices (left and right separately), we delineated four macro-anatomical landmarks: 1) the anterior Heschl’s Gyrus; 2) the superior temporal gyrus; 3) the superior temporal sulcus and 3) the middle temporal gyrus (see ***Figure 13*** and ***Figure 14***). These landmarks were used as additional information to determine the cost that is minimized during curvature based non-linear alignment in our tailored approach (i.e. CBA+). As for the standard surface alignment, CBA+ was performed across 4 spatial scales (from very smooth to slightly smooth curvature maps) which is shown to improve curvature based alignment overall (***Frost and Goebel, 2012; Tardif et al., 2015***). Both CBA and CBA+ were performed using dynamic group averaging. The surface alignment yielded a mapping between each individual to a group average brain, each consisting of the same number of vertices.

**Figure 14.**
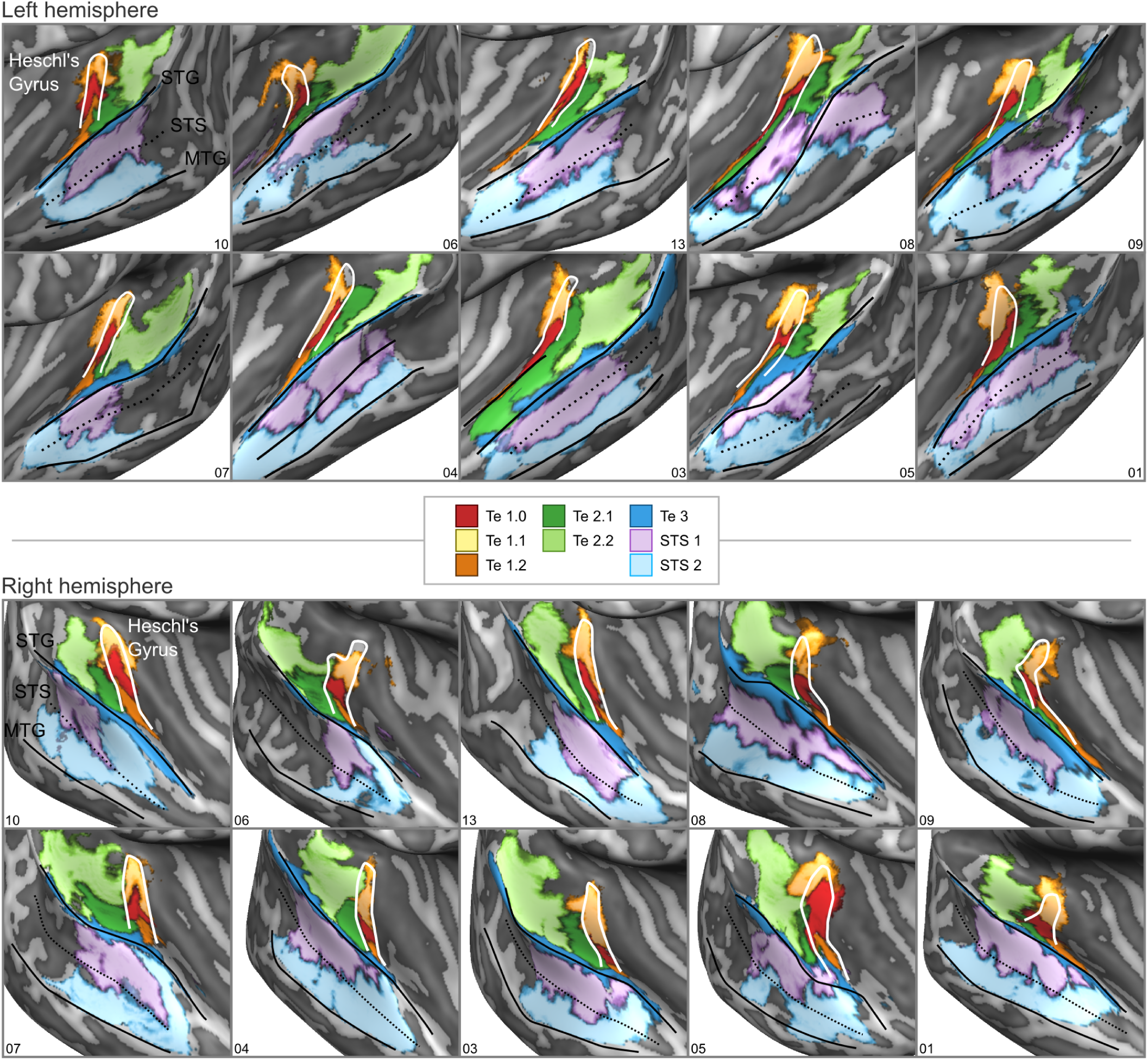
cyto-architectonic areas of ***Morosan et al. (2001***, 2005); ***Zachlod et al. (2020***) sampled on the inflated cortical surfaces for each individual brain in the post mortem dataset. Anterior Heschl’s Gyrus, superior temporal gyrus (STG), superior temporal sulcus (STS), and middle temporal gyrus (MTG) are indicated as line drawings.

To evaluate the effect of alignment, we computed the overlap across individuals for each of the cyto-architectonic areas. We compared our tailored alignment procedure to the original volumetric Colin27 alignment (***Evans et al., 2012***), spherical rigid body alignment, and non-linear standard cortex based alignment (see ***Figure 14***).

### In vivo data

#### MRI acquisition

We have used the dataset^2^ described in (***Sitek et al., 2019***). This dataset includes: (I) T_1_ weighted (T_1_w), proton density weighted (PDw) and 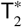 weighted 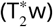 anatomical images collected (using a modified MPRAGE sequence) at a resolution of 0.7 mm isotropic (whole brain); (II) functional images at collected at a resolution of 1.1 mm isotropic (partial coverage, coronal-oblique slab, multi band factor=2; GRAPPA = 3) in response to the presentation of natural sounds (168 natural sounds; 24 runs divided in four cross validation splits of 18 training and 6 testing runs each (126 training sounds and 42 testing sounds per split).

#### Cortical segmentation and alignment

Segmentations of both the white matter - gray matter interface and outer gray matter (also called gray matter - cerebrospinal fluid interface) were done following BrainVoyager 2.8.4’s advanced segmentation pipeline (***Kemper et al., 2018***) and using the automatic bridge removal tool (***Kriegeskorte and Goebel, 2001***). Manual corrections were done in ITK-SNAP (***Yushkevich et al., 2006***). All follow up analyses were performed by sampling (anatomical and functional) data onto the middle gray matter surfaces (defined using the equidistant methods (***Waehnert et al., 2014; Kemper et al., 2018***) by the combination of inner and outer gray matter surfaces). This allowed us to minimize partial voluming with white matter, cerebrospinal fluid or superficial vessels. These surfaces can be seen for each individual in ***Figure 15***.

**Figure 15.**
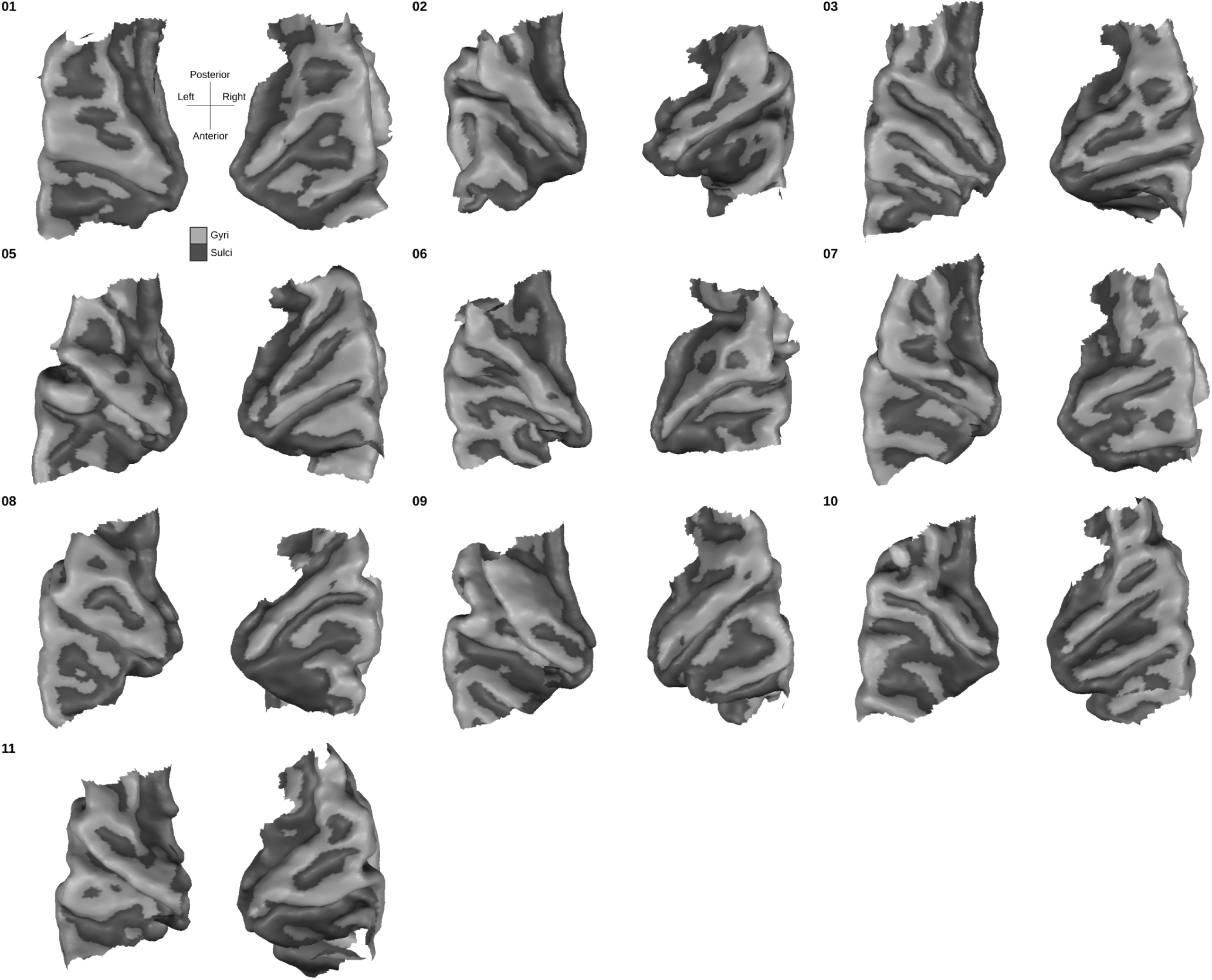
Individual superior temporal cortex middle-gray matter surface reconstructions from a bird’s eye (top-down) view. Dark gray colored indicate sulci and light gray indicates gyri.

The middle gray matter surfaces of all individuals were aligned using the procedure tailored to the superior temporal plane described above (CBA+). The resulting group average mesh from the in vivo dataset was aligned to the average post mortem mesh following the same procedure. This allowed us to overlay probabilistic cyto-architectonic areas onto the in vivo group average cortical surfaces and sample functional and anatomical data within each area.

#### Myelination maps

The processing steps followed to create myelination maps were similar to ***De Martino et al. (2015***). T_1_w images were divided by 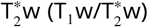 and the resulting division image was masked by the cortical gray matter segmentation. A histogram-based adaptive percentile threshold (based on iterative deceleration of percentile differences) on the 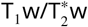 image was used to discard voxels with extreme intensities corresponding to vessels and regions in which the 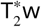 data were of insufficient quality. Maps were rescaled to range between 0-100. This step was necessary to match intensity ranges across subjects since we did not have quantitative measures. Values in the middle gray matter of the rescaled maps were sampled onto the surface mesh.

#### Tonotopy maps

The functional data were preprocessed using BrainvoyagerQX v2.8.4 (***Goebel, 2012***). Slice-scantime correction, motion correction, temporal high-pass filtering (GLM-Fourier, 6 sines/cosines) and temporal smoothing (Gaussian, kernel width of two acquisition volumes [i.e. 5.2 s]) were applied. Default options in BrainvoyagerQX v2.8.4 were used aside from the explicitly stated values. The functional images were then distortion corrected using the opposite phase encoding direction images using FSL-TOPUP (***Andersson et al., 2003***) as implemented in (***Smith et al., 2004***). The conversion between Brainvoyager file types to NIfTI, which was required to perform distortion correction, was done using Neuroelf version 1.1 (release candidate 2) ^3^ in Matlab version 2016a.

After pre-processing, functional images were transformed to Talairach space using BrainvoyagerQX v2.8.4 at a resolution of 1 mm isotropic. We estimated the voxels’ responses to each natural sound in a two step procedure (***Moerel et al., 2013; Santoro et al., 2014***). First, the hemodynamic response function (HRF) best characterizing the response of each voxel was obtained using a deconvolution GLM (with 9 stick predictors together with the noise regressors) on the training data (a subset of the functional runs). Second, the response to each natural sound (in training and test set runs separately per cross validation) was estimated with using a GLM analysis and the optimized HRF of each voxel. In addition to the predictors representing the experimental conditions (i.e. the individual stimuli), the analysis included noise regressors obtained using GLM-denoise (***Kay et al., 2013***). Note that the number of noise components and their spatial maps (allowing to derive the temporal regressors) where estimated on the training data only (i.e. separately per each cross-validation).

To estimate the voxels’ preference to the acoustic content (i.e. sound frequencies) we fitted (using Ridge Regression) the spectral sound representation obtained by passing the sounds through a cochlear filter model (128 logarithmically spaced filters, see (***Chi et al., 2005; Moerel et al., 2013***)) to the voxels’ responses (i.e. linearized encoding approach (***Kay et al., 2008***)). The frequency associated with the largest linear weight after fitting defined the preference of each voxel (see (***Moerel et al., 2012***) for more details on the procedure). Tonotopic maps were obtained by color coding (red to blue) the frequency preference (low to high) at each voxel.

## Acknowledgments

We thank Peer Herholz, Agustin Lage-Castellanos, and Fred Dick for their comments and advice at different stages of this project. The authors O.F.G. and F.D.M. were supported by NWO VIDI grant 864-13-012, and M.M. was supported by NWO VENI grant 451-15-012. The authors R.G and K.A. received funding from the Human Brain Project grant agreement no. 737691 (SGA2).

---

**Figure 10–Figure supplement 1.**
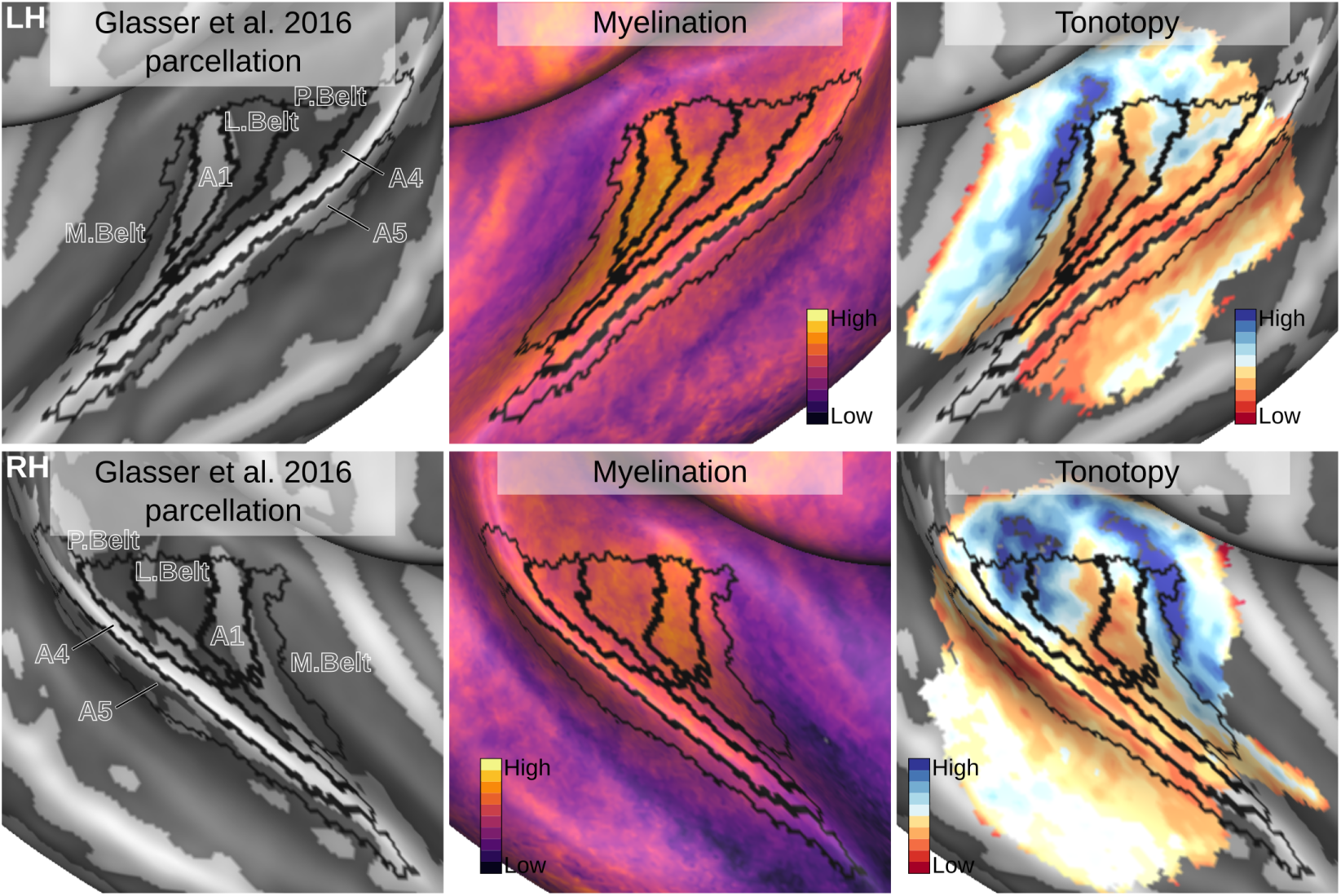
Relation between the in vivo MRI measures of the left hemisphere and the multi model MRI based parcellation. The multi modal MRI based parcellation from ***Glasser et al. (2016***) is delineated with black lines. The myelination index is computed from the division of T_1_w and 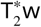 data. Tonotopy reflects the voxel-wise frequency preference estimated with fMRI encoding from the response to natural sound stimuli. All measures are sampled on the middle gray matter surfaces.

## Footnotes

1 The JuBrain atlas is available through the Atlas of the Human Brain Project https://jubrain.fz-juelich.de/

2 This dataset is available at: https://openneuro.org/datasets/ds001942/versions/1.2.0

3 http://neuroelf.net/

